# Liverwort MpSDR4 regulates abscisic acid sensitivity and gemma cup growth

**DOI:** 10.64898/2026.10.21.760454

**Authors:** Jiayi Shan, Jay Mie Kua, Janty Woojuh, Sean Cutler, Sean W. Graham, Liang Song

## Abstract

- Complex traits often evolve by co-opting pre-existing genetic components. Orthologs of the transcriptional co-factor SEED DORMANCY 4 (SDR4) are found across seed-bearing and seed-free land plants. In flowering plants, loss-of-function mutants of *sdr4* orthologs in rice and *Arabidopsis* exhibit opposite phenotypes in seed germination, a trait regulated by the phytohormone abscisic acid (ABA).
- We uncovered a major gene duplication event that closely preceded the origin of seed plants, by using a structure-informed molecular phylogenetic analysis. This duplication gave rise to the paralogous SDR4 and DIG/DIL clades, which are broadly retained across the seed plants.
- Investigating the role of SDR4 homologs in the common liverwort *Marchantia polymorpha*, a seed-free model organism with only a single SDR4-like homolog (MpSDR4), may provide insights into how SDR4 evolved to regulate dormancy and germination in other land plants. Loss-of-function mutants of Mp*sdr4* and transcriptome profiling showed that MpSDR4 modulates ABA sensitivity, consistent with the idea that SDR4 subsequently evolved to regulate hormone responses related to seed germination, with additional functional divergence between monocots and eudicots.
- Furthermore, Mp*sdr4* mutants exhibit elevated growth of gemma cups, contrasting the roles of SDR4 orthologs in asexual reproduction in liverwort with the regulation of seed traits in flowering plants.

## Introduction

Abscisic acid has played a crucial role in plant terrestrialization. Unlike fresh water algae, land plants need to cope with less predictable water availability. In bryophytes, ABA was co-opted as a ligand of the Pyrabactin Resistance 1-Like (PYL) receptor proteins (Sun *et al*., 2019). The ABA-dependent inhibition of protein phosphatase 2C (PP2C) activity allows the receptor-phosphatase-kinase regulatory module to sense environmental stresses through ABA. Evolution of receptor proteins have taken place independently in mosses and flowering plants to reduce the high background inhibition of PP2Cs, thus allowing more dynamic downstream signaling events in response to ABA (Zimran *et al*., 2025).

An evolutionarily conserved transcription factor ABSCISIC ACID INSENSITIVE 3 (ABI3) is essential for desiccation tolerance and other ABA responses in bryophytes (Khandelwal *et al*., 2010; Eklund *et al*., 2018). Orthologs of ABI3 and ABA are known to transactivate *Late Embryogenesis Abundant* (*LEA*) genes in spreading earthmoss *Physcomitrella patens*, common liverwort *M. polymorpha*, and many seed plants (McCarty *et al*., 1991; Marella *et al*., 2006; Eklund *et al*., 2018; Tian *et al*., 2020). In contrast to ABI3’s role in both spores and throughout the vegetative phase in *P. patens* and *M. polymorpha*, the function of ABI3 is most prominent during seed development and early stages of seedling establishment in flowering plants. Being one of the master regulators of seed development, ABI3 controls all major aspects of seed maturation (Gazzarrini & Song, 2024). Other evolutionarily conserved genes downstream of ABA signaling include *MOTHER of FT and TFL1* (*MFT*), a member in the phosphatidylethanolamine-binding (PEPB) family that is found in both seed-free and seed lineages (Hedman *et al*., 2009; Karlgren *et al*., 2011). Similar to ABI3, orthologs of MFT are known to regulate seed traits and germination in a wide range of monocot and eudicot species (Xi *et al*., 2010; Nakamura *et al*., 2011; Vaistij *et al*., 2018; Lu *et al*., 2023; Cai *et al*., 2023), suggesting these ABA-mediated regulation is broadly present in land plants. Compared to ABA’s inhibitory role in the phase transition from seed to seedlings, the hormone’s regulation of the vegetative-to-reproductive transition is less characterized. Lesions in the ABA biosynthesis gene *ABSCISIC ACID DEFICIENT2* (*ABA2*) and *ABI3* both result in earlier flowering (Kurup *et al*., 2000; Domagalska *et al*., 2010), corroborating the prevention of floral transition by ABA (Shu et al., 2018).

To obtain a comprehensive understanding of the regulatory network, it is highly important to functionally characterize novel regulators of ABA responses in addition to the canonical ABA signaling genes. SEED DORMANCY 4 (SDR4) is encoded by a quantitative trait locus that regulates seed dormancy in rice (Sugimoto *et al*., 2010). Its alleles regulate seed dormancy in response to precipitation and temperature in both the *japonica* and *indica* subspecies of rice (Zhao *et al*., 2022). These characteristics make SDR4 a promising candidate for grain improvement in cereals, for instance, to reduce pre-harvest sprouting. SDR4’s ortholog in Arabidopsis, *Arabidopsis thaliana SDR4-like* (*AtSDR4L*) and its paralogs, *Dynamic Influencer of Gene expression* (*DIGs*), and *DIG-likes* (*DILs*) are predominantly expressed in seeds, and are strongly responsive to abscisic acid (ABA) (Song *et al*., 2016; Gazzarrini & Song, 2024). Very intriguingly, mutant analysis within angiosperms has shown that the SDR4 homologs in *Arabidopsis* exhibit a seemingly opposite function to what is seen in rice SDR4: loss-of-function mutation of *SDR4* orthologs in rice and in Arabidopsis results in elevated and reduced seed germination, respectively (Cao *et al*., 2020; Zhao *et al*., 2022). In addition to the regulation of germination, AtSDR4L represses master transcription factors of seed development, and mutant seedlings exhibit stunted growth and overaccumulation of seed storage lipids (Zheng *et al*., 2022; Wu *et al*., 2022). These observations make it particularly interesting to broadly characterize the relationship in plants between SDR4 and abscisic acid (ABA), a plant hormone with a critical role in regulating seed dormancy and germination.

To better understand how evolution shapes SDR4-mediated regulation of seed traits, we investigated the phylogenetic history of the SDR4/DIG family using structure-informed alignments, and functionally characterized the SDR4 ortholog in *M. polymorpha*. We uncovered a major gene duplication event of *SDR4* that occurred around the origin of seed plants that gave rise to the paralogous *SDR4* and *DIG*/*DIL* clades. In *Marchantia*, MpSDR4 maintains sensitivity to ABA and related abiotic stresses. Additionally, MpSDR4 inhibits gemma cup formation and growth, possibly through an ABA-independent mechanism. These findings in extant liverwort and knowledge from *Arabidopsis* and rice strongly suggest an evolutionarily deep connection between the SDR4 family and ABA responses may have formed in early land plants.

## Methods and materials

### Identification of SDR4 homologs

Full-length amino acid sequence of *A. thaliana* SDR4 and its six paralogs, Dynamic Influencer of Gene expression 1 (DIG1), DIG2, and DIG-likes (DILs; DIL1 - 4) were used as query sequences for a preliminary BLASTp search across species of green plants. We further selected several putative SDR4 protein homologs from the preliminary BLASTp search for protein secondary structure prediction using AlphaFold Server (Abramson *et al*., 2024). *Klebsormidium nitens* (kfl00215_0250; a green alga), and representative land plants, including *Marchantia polymorpha* (Mp2g17570; a liverwort and a major subject of this thesis), *Selaginella moellendorffii* (437663; a lycophyte), *Adiantum capillus-veneris* (Adc31130; a fern), *Ginkgo biloba* (evm.model.chr2.251; a gymnosperm) and *A. thaliana* (AT1G27461 and AT3G48510; an angiosperm) were chosen for a thorough sampling across green lineages. The C-terminal protein fold containing 7-8 beta sheets alternating with several alpha-helices of these proteins, with slight variations across species, was superimposed using the matchmaker command supported by UCSF ChimeraX (Meng *et al*., 2023) to confirm the structure conservation.

The protein sequences of SDR4 protein homologs from these six species that encode the conserved C-terminal protein fold were used as queries in BLASTp searches against various plant genome databases or datasets (Table S1). For the databases with an online BLASTp server, the default BLASTp parameters were used, and all the unique hits were selected for further structural examination using the AlphaFold Server (Abramson *et al*., 2024). When genomes of interest were not available on online plant genome databases, genome annotations with the peptide sequences deposited by genome assembly papers were downloaded for a local BLASTp search (Camacho *et al*., 2009). Local BLAST databases were created using the command line “*makeblastdb -in protein.fa -dbtype prot -out db/protein*”, followed by BLASTp search. All hits with an E-value smaller than 0.001 were chosen for further protein secondary structure verification. After compiling all the hits from various species, the presence of the structurally conserved C-terminal protein fold were checked using AlphaFold Server prediction (Abramson *et al*., 2024), followed by superimposition on conserved C-terminal protein fold shown in Figure 1b using UCSF ChimeraX (Meng *et al*., 2023); the sequence information is detailed in Table S1. Sequences without specific notes in Table S1 show a well-superimposed C-terminal protein fold; these homologs were included in multi-sequence alignments.

**Figure 1.**
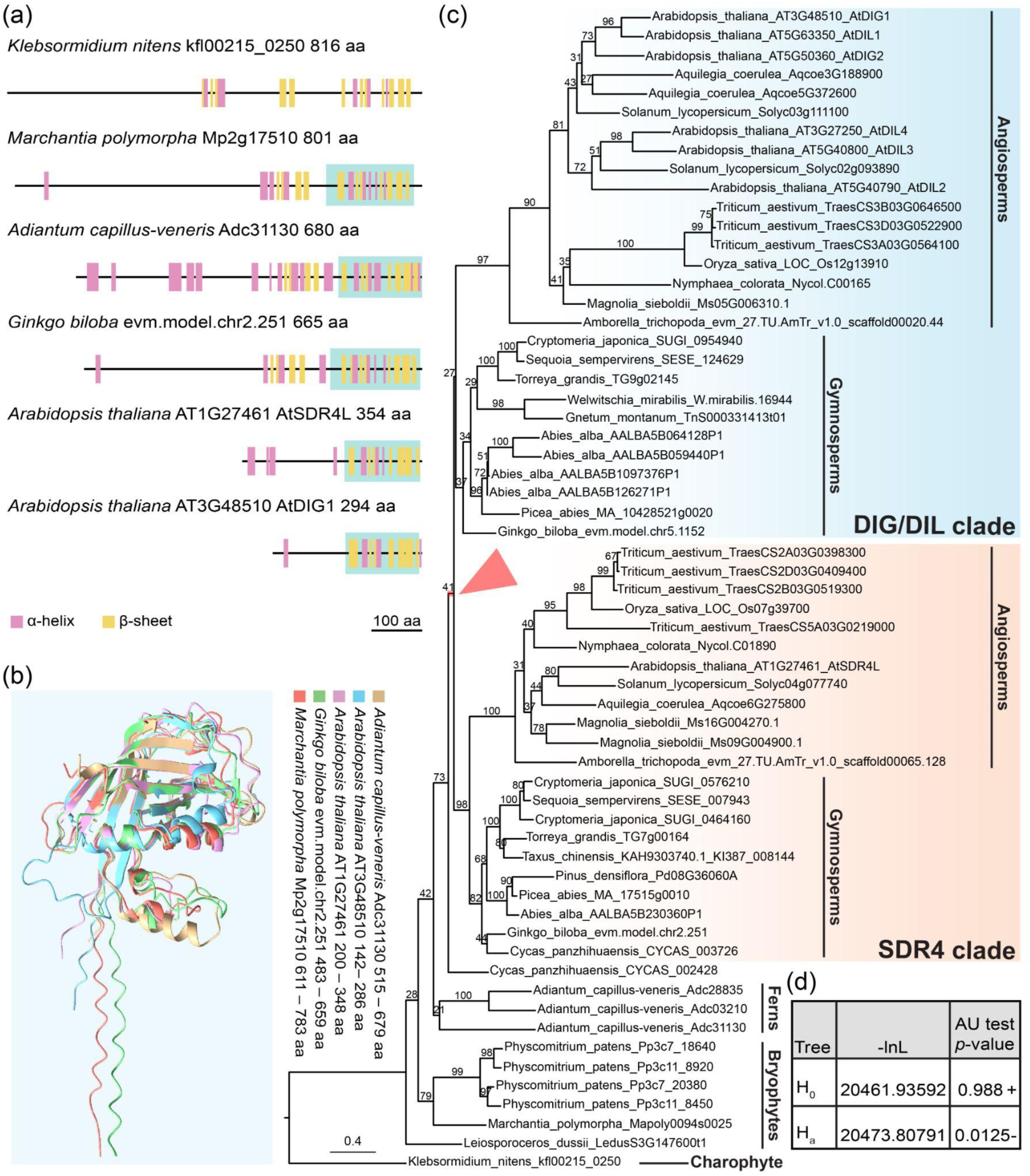
Structure and phylogeny of the SDR4 protein family. (a) Structural prediction of full-length SDR4 protein homologs across green lineages by AlphaFold 3. The conserved C-terminal protein fold shown in (b) is shaded in teal. (b) Superimposition of the C-terminal protein fold of the five SDR4 protein homologs from the land plants shown in (a). (c) Phylogram of the SEED DORMANCY 4 (SDR4) protein family inferred from a maximum likelihood analysis of SDR4 protein sequences from 24 plant species representing major lineages in streptophytes. The red-highlighted branch marked by the arrowhead indicates where a major gene duplication event that led to the SDR4 clade and DIG/DIL clade may have occurred. Bootstrap support values are indicated on branches. The scale bar shows the estimated number of substitutions per site. *M. polymorpha* SDR4 ortholog Mapoly0094s0025 corresponds to Mp2g17570. (d) Results of an approximately unbiased (AU) test comparing the best tree (H_0_), shown in (c), with an alternative hypothesis (H_a_) in which the angiosperm SDR4 and DIG/DIL are constrained to form a monophyletic group. -lnL: negative log-likelihood score; a “+” sign represents the tree is within the 95% confidence sets; “-” indicates significant rejection of the alternative topology. See also Figure S2 and Table S1.

### Multiple sequence alignment (MSA) and trimming

A multi-sequence alignment of amino acid sequences of the selected SDR4 homologs was generated using MUSCLE v3.8.31(Edgar, 2004) implemented in Aliview (Larsson, 2014) using default parameters. The resulting alignment was automatedly trimmed by trimAl (Capella-Gutiérrez *et al*., 2009) using the following command line: “*trimal -in inputfile.fas -out outputfile.fas -automated1*”.

### Generation of maximum likelihood (ML) tree and hypothesis testing

Maximum-likelihood (ML) analysis was performed using IQ-TREE 2 (Minh *et al*., 2020) multicore version 2.4.0. The best-fit protein model (JTT+G4) was chosen based on the Bayesian information criterion (BIC) using ModelFinder (Kalyaanamoorthy *et al*., 2017). Branch support for the resulting tree was assessed using the standard nonparametric bootstrap with 100 replicates.(Felsenstein, 1985) The final ML tree was visualized using FigTree v1.4.4 (Rambaut, Andrew, 2018).

The constrained ML tree search and the subsequent hypothesis testing using approximately unbiased (AU) test (Shimodaira, 2002) were carried out by IQ-TREE multicore version 2.3.6 (Minh *et al*., 2020). The monophyly of the angiosperms SDR4 and angiosperms DIG/DIL was constrained. The internal relationships within the constrained monophyletic group and the relationships among the remaining taxa were not specified. The constrained ML tree was generated and compared with the unconstrained ML tree under the JTT+G4 model. Bootstrap with 10000 replicates were performed using the RELL method (Kishino *et al*., 1990) for the AU test (Shimodaira, 2002).

### Plant materials and growth conditions

*Marchantia polymorpha L.* subsp. *ruderalis* accessions Takaragaike 1 (Tak-1) and Takaragaike 2 (Tak-2) were used as the wild-type male and female, respectively (Ishizaki *et al*., 2008). *Marchantia polymorpha* plants were cultured on half-strength Gamborg’s B5 medium (Gamborg *et al*., 1968) supplemented with 1% (w/v) agar (PhytoTech Labs, Cat # A111) under either continuous white light at 22°C or in long-day conditions (16 hours light at 22°C and 8 hours dark at 20°C).

### Plasmid construction

Loss-of-function mutants of Mp*SDR4* (Mp2g17570) were generated with the CRISPR/Cas9 system using the OpenPlant toolkit and Golden Gate Loop Assembly as previously described (Sauret-Güeto *et al*., 2020). Two sets of gRNAs (gRNA1&3 and gRNA2&4 were designed using CRISPRdirect (Naito *et al*., 2015) to target the first exon of Mp*SDR4* (Figure 2a). Guide RNA1 & 3 (or gRNA2 & 4) was cloned into L1_lacZgRNA-Ck2 (Addgene, Cat # 136137) and L1_lacZgRNA-Ck3 (Addgene, Cat # 136136), respectively, using primers listed in Table S2 and BpiI (Thermo Fisher Scientific, Cat # ER1011) for Type IIS assembly. Resulting L1_gRNA1 (or gRNA2)-Ck2 and L1_gRNA3 (or gRNA4)-Ck3 were then assembled with L1_HyR-Ck1 (Addgene, Cat # 136125) and L1_Cas9-Ck4 (Addgene, Cat # 136135) were then assembled into the designated vector pCsA (Addgene, Cat # 136067) using loop assembly by SapI (New England Biolabs, Cat # R0569S). Assembled plasmids were transformed into *E.coli* strain 10β (New England Biolabs, Cat # C3019H), selected by blue-white screening on plates supplemented with 1 mM IPTG (Gold Biotechnology, Cat # I2481C), 40 μg/mL X-Gal (Rose Scientific Ltd, Cat # 7240-90-6), and 50 μg/mL spectinomycin (Gold Biotechnology, Cat # S-140-25), and extracted plasmids were validated by Sanger sequencing.

**Figure 2.**
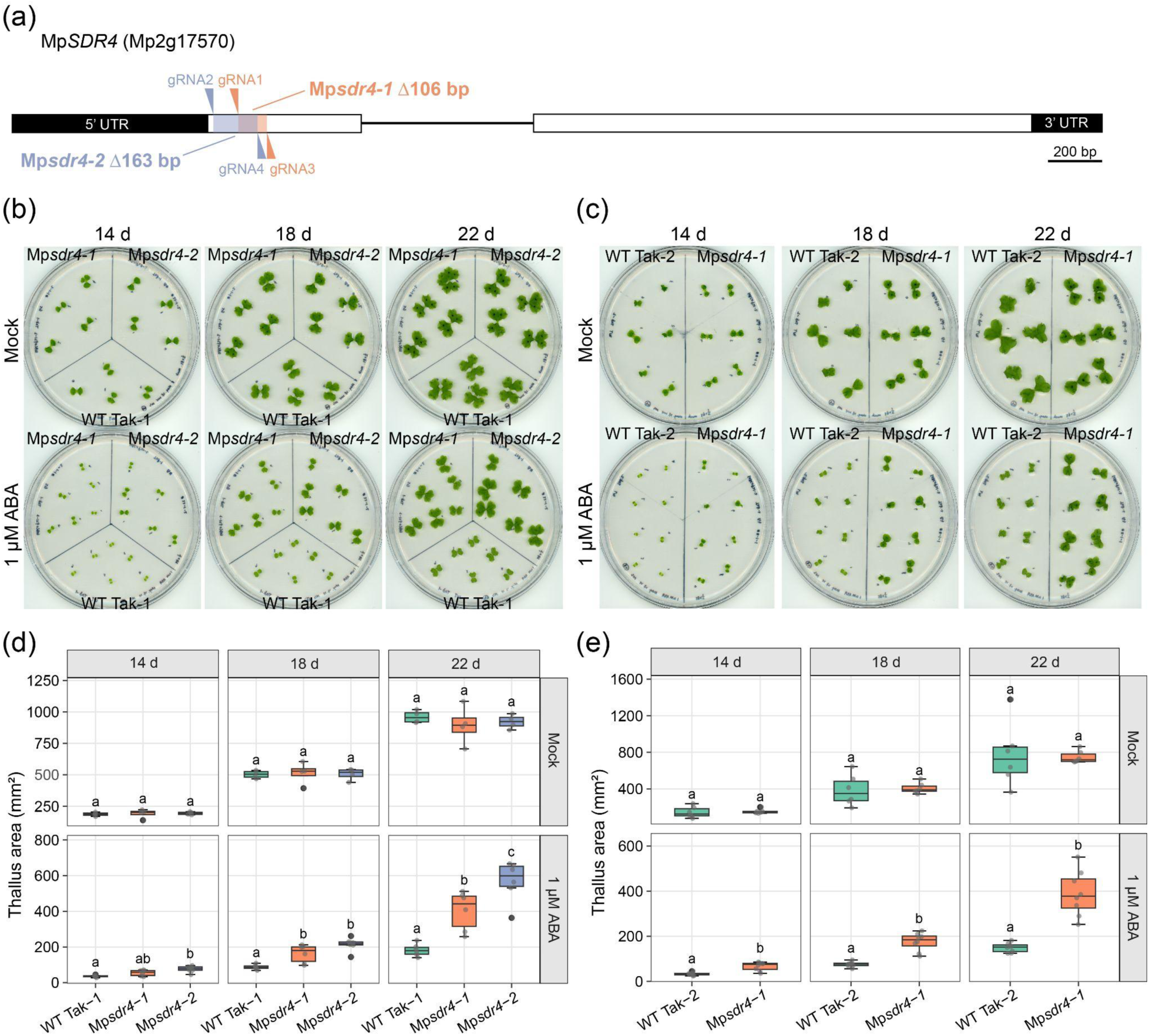
MpSDR4 positively regulates ABA sensitivity. (a) Schematic representation of MpSDR4 gene model, and mutant alleles generated by CRISPR/Cas9. Arrowheads indicate the gRNA target sites. The deletion in two independent alleles was colored in purple and orange, respectively. Untranslated regions are shown as black boxes, exons as white boxes, and the intron as a line connecting exons. (b - c) 14-, 18-, and 22-day-old Mp*sdr4* mutants and wild-type plants in the Tak-1 (b) and Tak-2 (c) backgrounds, grown under long-day conditions on medium without (mock) or with supplemented ABA. (d - e) Thallus area of Mp*sdr4* mutants and wild-type plants, shown in (b) and (c), respectively. Different letters represent groups with significant differences (n=4 plants for mock and n=6 plants for 1 µM ABA treatment, *p* < 0.05, one-way ANOVA followed by Tukey’s HSD test; n=6 plants for mock and n=8 plants for 1 µM ABA treatment, *p* < 0.05, two-tailed Student’s *t* test). See also Figure S3.

### Agrobacterium-mediated transformation using regenerating thalli

The final constructs were transformed into *Agrobacterium tumefaciens* strain GV3101. Regenerating thalli were prepared as previously described (Kubota *et al*., 2013). Approximately 60 - 70 regenerating thalli per construct were inoculated with 20 mL half-strength Gamborg’s B5 liquid medium (PhytoTech Labs, Cat # G768) containing 1% sucrose, 100 μM acetosyringone (Cayman Chemical Co., Cat # 23224), and 1 mL *Agrobacterium*-suspension (OD_600_ around 2) for 30 mins. The plant pieces were then desiccated on sterile filter papers under a laminar flow hood for 30 minutes. After desiccation, the plant tissues were transferred to co-culture media composed of half-strength Gamborg’s B5 medium supplemented with 1% sucrose and 100 μM acetosyringone for three days in the dark. Following co-cultivation, the thalli pieces were transferred to half-strength Gamborg’s B5 media, supplemented with 300 μg/mL Timentin (Gold Biotechnology, Cat # T-104-10), and cultured for three days under continuous light. Thalli were then plated on half-strength Gamborg’s B5 media supplemented with 10 μg/mL Hygromycin B (Gold Biotechnology, Cat # H-270-10) and 300 μg/mL Timentin for selection. Successful transformants appeared after three weeks.

### Identification of CRISPR mutants

To assess genome editing events, gemmae derived from regenerating transgenic T1 plants (i.e., the asexual G1 generation) were examined by polymerase chain reaction (PCR) amplification followed by Sanger sequencing. Small pieces (3 × 3 mm) of thalli from each plant were placed in 1.5 mL microcentrifuge tubes containing two metal beads and 100 μL genotyping buffer (100 mM Tris-HCl, 1 M KCl, and 10 mM EDTA, pH 9.5). Plant tissues were homogenized using a TissueLyser (30 sec at 30 beats/sec) followed by incubation at 80 °C for 5 mins. The resulting DNA was diluted by adding 400 μL of water before PCR using primers listed in Table S2. Stable-editing events were further confirmed in the G2 and G3 generations, and plant materials from the G2 generation and beyond were used for downstream experiments.

### Abscisic acid (ABA) and antabactin (ANT) treatment

Experiments were performed under continuous white light at 22°C or under long-day conditions. Abscisic acid (Gold Biotechnology, Cat # A-050-1) was dissolved in 100% (v/v) ethanol to make the 100 mM ABA stock solution and followed by a serial dilution with sterile water. Diluted ABA solution was added to 50 mL half-strength Gamborg’s B5 medium supplemented with 1% (w/v) agar to make 1 μM, 5 μM and 10 μM ABA treatment plates. The mock treatment used ethanol in place of ABA. Antabactin (Vaidya *et al*., 2021) was dissolved in 100% DMSO to make a 40 mM stock solution. For ANT treatment, the final concentration of ANT was diluted to 10 μM in 50 mL half-strength Gamborg’s B5 medium supplemented with 1% (w/v) agar. For ANT and ABA treatment, the final concentration of ANT was diluted to 10 μM and ABA was diluted to 1 μM in 50 mL half-strength Gamborg’s B5 medium solidified with 1% (w/v) agar. The mock treatment used DMSO in place of ANT.

### Osmotic and salt stress treatment

Experiments were performed under either continuous white light at 22°C or under long-day conditions. For osmotic stress treatment, gemmae of Tak-1 wild type, *Mpsdr4-1* and *Mpsdr4-2* mutants in Tak-1 background were plated on half-strength Gamborg’s B5 medium solidified with 1% (w/v) agar supplemented with D-mannitol (Alfa Aesar, Cat # A14030) at 100 mM and 150 mM. Half-strength Gamborg’s B5 medium supplemented with 1% (w/v) agar was used as mock treatment. The experiment was validated using wild type and Mp*sdr4-1* mutant in the Tak-2 background.

For salt stress treatment, gemmae of Tak-1 wild type, Mp*sdr4-1* and Mp*sdr4-2* mutants in Tak-1 background were plated on half-strength Gamborg’s B5 medium supplemented with 1% (w/v) agar and 60 mM sodium chloride (Thermo Fisher Scientific, Cat # BP358-212). Half-strength Gamborg’s B5 medium solidified with 1% (w/v) agar was used as mock treatment. The experiment was repeated using wild type and Mp*sdr4-1* mutant in the Tak-2 background.

### RNA extraction for quantitative RT-PCR analysis and mRNA library preparation

Total RNA was extracted using the Spectrum™ Plant Total RNA kit (MilliporeSigma, Cat# STRN50). RNA was then subjected to 2 U DNase, followed by 25 minutes of incubation at 37°C using TURBO DNA-free™ Kit (Thermo Fisher Scientific, Cat # AM1907). For RT-qPCR, the DNase-treated RNA was reverse transcribed using Maxima H Minus Reverse Transcriptase (Thermo Fisher Scientific, Cat # EP0752) at 50°C for 30 mins and 85°C for 5 mins. qPCR was performed using PowerUp™ SYBR™ Green Master Mix (Thermo Fisher Scientific, Cat # A25742) in a QuantStudio™ 3 System. *APT* (Mp3g25140) was used as an internal control. Primer sequences are listed in Table S2. Significance differences (*p* < 0.05) were tested using one-tailed Student’s *t* test. For library preparation, 350 ng of total RNA was subjected to 2 U DNase, followed by incubation at 37°C for 25 minutes using TURBO DNA-free™ Kit (Thermo Fisher Scientific, Cat # AM1907) before NEBNext Ultra™ II Directional RNA Library Prep Kit for Illumina (New England Biolabs, Cat # E7760S) according to the manufacturer’s protocol.

### RNA-seq analysis

Libraries were paired-end (PE150) sequenced on NovaSeq X Plus Series. Raw sequencing reads were trimmed for adaptors using fastp v0.12.4 with default parameters (Chen *et al*., 2018). Then, the filtered reads were aligned to the *Marchantia polymorpha* MpTak_v7.1 standard genome (Tanizawa *et al*., 2025) and annotation using STAR 2.7.9a with the arguments “*--runMode alignReads --outFilterMultimapNmax 10 --outFilterMismatchNoverLmax 0.05 --quantMode GeneCounts --outSAMtype BAM SortedByCoordinate --outReadsUnmapped Fastx*” (Dobin *et al*., 2013). To visualize mapped reads, BAM files were converted to TDF files using igvtools v2.5.3 “count” command with arguments “--includeDuplicates -w 10 -z 5”. and loaded into the Integrative Genomics Viewer v2.17.4 (Robinson *et al*., 2011) along with the MpTak_v7.1 standard genome.

To identify differentially expressed genes, read count was extracted from the fourth column of each “*ReadsPerGene.out.tab*” file and merged to create a single read count matrix for all ten library samples. The lowly expressed genes were filtered by counts per million (CPM) values greater than 1 in at least 20% of the samples and the filtered raw reads were normalized by Trimmed Means of M-values (TMM) using edgeR package v4.2.2 (Robinson *et al*., 2010) in RStudio v2024.04.2+764 (RStudio Team, 2020). Differential expression analyses were conducted using *limma* package v3.60.6 (Ritchie *et al*., 2015) with the filtered TMM-normalized reads using voom transformation followed by usual limma pipelines. The principal component analysis using multidimensional scaling was performed on the ten library samples using filtered normalized reads with the “*plotMDS*” function. Differentially expressed genes (DEGs) were identified by comparing the response of wild-type Tak-1 and Mp*sdr4-2* under 150 mM D-mannitol treatment, and by assessing the response of wild-type Tak-1 and Mp*sdr4-2* to 150 mM D-mannitol treatment versus mock treatment. Genes exhibiting Log2 fold changes greater than 0.585 or less than -0.585 with an FDR-adjusted p-values below 0.05 were considered significantly up- or down-regulated, which were used for generating the Venn diagram. A union of genes that are significantly up-regulated in wild-type Tak-1 and Mp*sdr4-2* under 150 mM D-mannitol treatment were used to filter through all detected genes after CPM filtering of these two datasets for generating the scatterplot.

Motif analysis was carried out using the MEME suite web server. DNA sequences 500 bp upstream of the transcription start site (TSS) of each gene were extracted according to the GFF file of MpTak_v7.1 standard genome using package *Biostrings* v2.72.1 (H. Pagès, 2017) and package *GenomicRanges* v1.56.2 (Lawrence *et al*., 2013) in RStudio v2024.04.2+764. Analysis of Motif Enrichment (AME) v5.5.85.5.8 (McLeay & Bailey, 2010) and *de novo* motif discovery with STREME v5.5.8 (Bailey, 2021) were conducted for DEGs with default parameters using 500 bp sequence upstream of TSS of all genes in *Marchantia polymorpha* as control. The Arabidopsis DAP-seq motif (O’Malley *et al*., 2016) was selected as the input motif database for AME.

### Imaging and phenotypic measurement

The gemma cup images were captured using Leica M205 FCA Fluorescence stereo microscope with objective lens and magnification at 3.2. The area of thallus and diameter of gemma cups were measured using Fiji(Schneider *et al*., 2012) based on plate scanning using Epson Perfection V600 Photo at resolution 1200 dpi with Tag Image File Format (TIFF) image type. Each individual plant was treated as a biological replicate. For the thallus area, thresholding was adjusted to encompass the thallus. The wand tracing tool was used to select the area and the function “measure” was used to quantify the defined area. For gemmae cup diameter, each visible cup of an individual plant was measured three times to calculate the average length. The measurements of diameter across multiple cups of an individual plant were then averaged to obtain the mean diameter for each biological replicate. For plants that have a single cup, the mean cup diameter of the plant is based on the measurements of the single cup. For plants without any visible cups, the mean diameter was set at “0”. Final average was calculated across the diameter of all biological replicates. Statistical analyses were carried out in R version 4.4.0 (R Development Core Team). Significance differences (*p* < 0.05) were tested using two-tailed Student’s *t* test or one-way ANOVA with the Tukey’s honestly significant difference (HSD). Sample sizes and specific statistical methods used are provided in figure legends.

### Accession numbers

Gene nomenclature is obtained from MarpolBase (https://marchantia.info/): Mp*SDR4* (Mp2g17570; JGI 3.1 sequence ID: Mapoly0094s0025), Mp*LEA1* (Mp5g23710), Mp*DHN1* (Mp6g07540), Mp*CGAM1* (Mp6g04830), Mp*APT* (Mp3g25140), Mp*NCED* (Mp2g07800), Mp*MFT* (Mp2g12650), Mp*ABI3A* (Mp5g08310), Mp*LEAL1* (Mp4g09300), Mp*LEAL2* (Mp4g14860), Mp*LAXR* (Mp5g06970), Mp*EXP4* (Mp1g22730), and Mp*PL1.3* (Mp3g01000).

## Results

### A major *SDR4* gene duplication closely preceded with the origin of seed plants

To explore the evolutionary history of SDR4 across green lineages, we performed a search of SDR4 protein homologs from 24 representative species of Streptophyta (Table S1). AlphaFold 3 (Abramson *et al*., 2024) predicted that these proteins feature an N-terminal disordered region and a structurally conserved C-terminal protein fold containing 7-8 β-sheets, alternating with several ɑ-helices (Figure 1a). AlphaFold 3 also predicted the C-terminal region of SDR4 homologs from representative seed-free and seed-bearing plants all fold to be a similar β-barrel-like structure (Figure 1b). PANTHER (Thomas *et al*., 2022) search classified candidate SDR4 homologs as von Willebrand factor A domain proteins, typical functions of which include ligand binding and protein– protein interactions (Whittaker & Hynes, 2002).

We used the conserved C-terminal protein fold from selected species to retrieve SDR4 protein homologs across major land-plant clades and generated a structure-informed multiple species alignment (MSA) (Figure S1) and maximum-likelihood (ML) tree inference (Figure 1c). As expected, the resulting tree topology shows that the seed plants are nested among the other seed-free land plants. The optimal topology implies that a major gene duplication led to two distinct seed-plant SDR4-like clades (Figure 1c), each containing nearly full complements of the major lineages of gymnosperms and angiosperms. We label these seed-plant clades as SDR4 or DIG/DIL here, as each also contains well-supported SDR4 and DIG/DIL angiosperm clades (with 100% and 97% bootstrap support, respectively). Given this tree topology, a major gene duplication event must have closely preceded the origin of the seed plants, either on the branch highlighted in red or the one immediately preceding it (Figure 1c); we suspect that a single *Cycas panzhihuaensis* sequence recovered as a sister group to the seed-plant SDR4 and DIG/DIL clades here represents a weakly misinferred placement of it from the DIG/DIL seed-plant clade, which otherwise lacks cycads (note that the intervening branches are poorly supported: 41% and 27%, respectively). Still, the broad seed-plant (spermatophyte) SDR4 clade is well supported (98%), but the branch leading to the DIG/DIL seed-plant clade (ignoring the likely misplaced Cycas sequence) is not. We therefore performed a constrained tree search (Minh *et al*., 2020), imposing an alternative sister-group relationship between the angiosperm SDR4 and DIG/DIL sequences (Figure S2), in an arrangement that would instead imply that a different duplication event gave rise to the paralogous SDR4 clade and DIG/DIL angiosperm clades (i.e., a much later gene duplication near the origin of angiosperms, with other unmarked duplications then implied in gymnosperms). However, the approximately unbiased (AU) test (Shimodaira, 2002) rules out this constrained alternative hypothesis (*p* < 0.05), supporting the unconstrained ML tree (Figure 1c) as the best-scoring topology (Figure 1d), which in turn validates our inference that there was a single major gene duplication event near the origin of seed plants that resulted in the SDR4 and DIG/DIL seed-plant clades (Figure 1c).

### MpSDR4 modulates sensitivity to abscisic acid and osmotic stresses in

#### Marchantia

Bryophytes hold a key phylogenetic position in land plants, with the extant clades possibly representing the sister group of extant vascular plants (One Thousand Plant Transcriptomes Initiative, 2019), and so they should be considered for a broad understanding of SDR4 function and evolution. Moreover, *M. polymorpha* possesses a genetically tractable system for functional studies (Ishizaki *et al*., 2016). Based on the phylogenetic analysis, we identified a single-copy ortholog of *SDR4* in *M. polymorpha*, Mp2g17570, which we designated as Mp*SDR4*. To investigate the functions of Mp*SDR4*, we employed CRISPR/Cas9-mediated mutagenesis and generated two independent alleles in the male Tak-1 and female Tak-2 backgrounds. Transformants expressing guide RNA (gRNA) 1 and gRNA 3 resulted in an identical deletion of 106 bp in Tak-1 and Tak-2, which we named *Mpsdr4-1*. A transformant expressing gRNA 2 and gRNA 4 resulted in a deletion of 163 bp in the Tak-1 background, which we named *Mpsdr4-2* (Figure 2a and S3a). These deletions caused a frameshift followed by a premature stop codon in both mutant alleles: Mp*sdr4-1* encodes an altered protein of 96 amino acids (aa), while Mp*sdr4-2* produces a protein consisting of only 15 aa. Compared to the 801-aa wild-type MpSDR4, the severely truncated mutant alleles are highly likely to be nonfunctional (Figure S3a). Because mutants of SDR4 orthologs in rice and *Arabidopsis* exhibit germination-related phenotypes and altered sensitivity to ABA, we evaluated the ABA sensitivity of Mp*sdr4* mutants using a gradient of ABA concentrations. Exogenous ABA inhibits the growth of both wild-type and mutant plants in a dose-dependent manner (Figure S3b). Additionally, Mp*sdr4* mutants in both Tak-1 and Tak-2 backgrounds grow significantly larger (*p* < 0.05, one-way ANOVA with Tukey’s HSD test for Tak-1, and two-tailed Student’s *t* test for Tak-2) than wild-type plants under ABA treatment (Figure 2b-2e and S3b-S3d). Mp*sdr4* mutants exhibit reduced sensitivity to ABA under both a long-day lighting scheme (Figure 2) and continuous illumination (Figure S3). These results indicate that MpSDR4 helps maintain ABA sensitivity during vegetative growth in *Marchantia*.

ABA has a major and evolutionarily conserved role in mediating responses to osmotic stress (Komatsu *et al*., 2020). Because Mp*sdr4* mutants show reduced sensitivity to exogenous ABA, we predicted that mutant plants should also be less sensitive to osmotic stress induced by D-mannitol and sodium chloride. Consistent with this prediction, Mp*sdr4* mutants in Tak-1 background grow significantly larger (*p* < 0.05, one-way ANOVA with Tukey’s HSD test) than wild type on plates supplemented with 100 mM and 150 mM D-mannitol (Figure 3a and 3b). The effect is similar but subtler in Tak-2 background, where we observed a significant size difference (*p* < 0.05, two-tailed Student’s *t* test) at a later developmental stage on medium supplemented with the higher concentration (150 mM) of D-mannitol (Figure S4a and S4b). Corroborating the physiological differences, two ABA-responsive genes, *LATE EMBRYOGENESIS ABUNDANT1* (*LEA1*) and *DEHYDRIN1* (*DHN1*), express to a significantly lower level (*p* < 0.05, one-tailed Student’s *t* test) in Mp*sdr4* mutant than in wild-type control on plates supplemented with 150 mM D-mannitol (Figure 3c). Similarly, Mp*sdr4* mutants in both Tak-1 and Tak-2 backgrounds are significantly larger (*p* < 0.05, one-way ANOVA with Tukey’s HSD test for Tak-1, and two-tailed Student’s *t* test for Tak-2) than corresponding wild-type controls on plates supplemented with 60 mM sodium chloride (Figure 3d and 3e). Collectively, these results show that vegetative growth of Mp*sdr4* mutants is less sensitive to osmostress and salt, supporting the idea that MpSDR4 attenuates ABA responses elicited by abiotic stresses.

**Figure 3.**
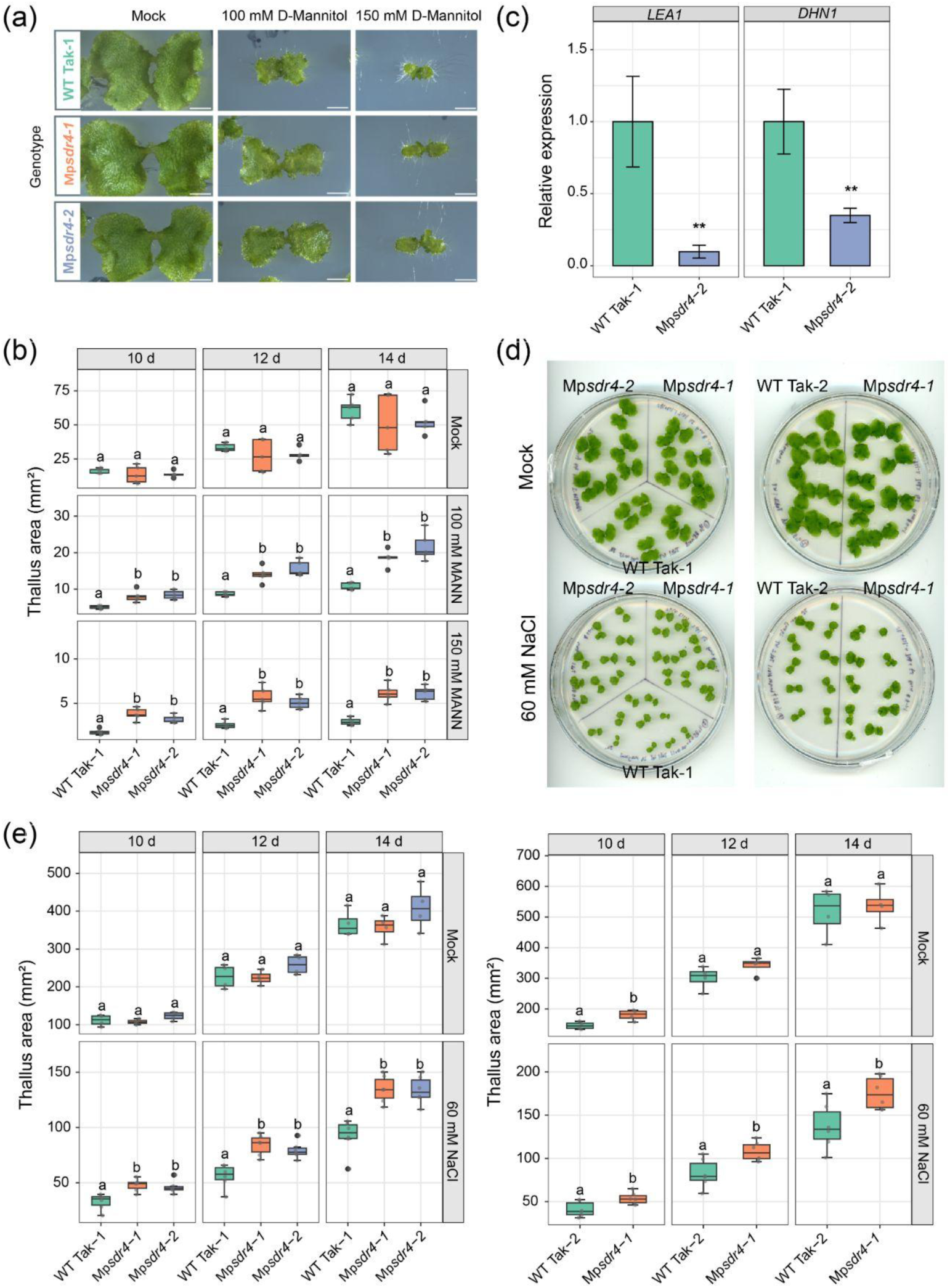
Mp*sdr4* mutants are hyposensitive to osmostress. (a) 17-day-old plants grown on medium under long-day conditions, without (mock) and with 100 mM and 150 mM of supplemented D-mannitol medium. Scale bars = 5 mm. (b) Thallus area of Tak-1 wild type and Mp*sdr4* mutants measured at 2-day intervals from day 10 to day 14. Plants were grown under long-day conditions on mock medium and medium containing indicated concentration of D-mannitol. Different letters represent significant differences (n=5, *P*<0.05, one-way ANOVA followed by Tukey’s HSD test). (c) Relative expression level of *LEA1* and *DHN1* in *Mpsdr4-2* compared to wild-type Tak-1 grown on medium supplemented with 150mM D-mannitol. Transcript levels were normalized to *ADENINE PHOSPHORIBOSYL TRANSFERASE* (Mp*APT*). Data are presented as mean ± SE. Significant differences are indicated by asterisks (\**p* < 0.05, \*\**p* < 0.01, one-tailed Student’s *t* test). (d) 14-day-old wild-type Tak-1 and Mp*sdr4* mutants (left), and wild-type Tak-2 and Mp*sdr4* mutant (right) grown on mock medium and medium containing 60 mM NaCl . (e) Thallus area of wild-type Tak-1 and *Mpsdr4* mutants (left), and wild-type Tak-2 and *Mpsdr4* mutants (right) measured at 2-day intervals from day 10 to day 14. Plants were grown under long-day conditions on mock medium and medium containing 60 mM NaCl treatment. Different letters represent significant differences (n = 4 for mock and n = 6 for NaCl treatment, *p* < 0.05, one-way ANOVA followed by Tukey’s HSD test for Tak-1, and two-tailed Student’s *t* test for Tak-2). See also Figure S4.

### Transcriptome analysis support the attenuation of ABA signaling in response to osmotic stress in Mp*sdr4* mutants

To systematically examine the regulation of osmotic responses by MpSDR4, we carried out RNA-seq using 20 DAS wild-type Tak-1 and Mp*sdr4-2* mutants grown on either mock medium or medium supplemented with 150 mM D-mannitol. We observed that transcriptomes are mainly influenced by the mannitol treatment and genotype (Figure 4a). When comparing mannitol-treated samples with mock-treated samples of the same genotype, wild-type Tak-1 and Mpsdr*4-2* mutant have 7,509 (Tak-1) (3,676 up, 3,833 down, p.adjust < 0.05) and 6,305 (sdr4-2) (2,927 up, 3,378 down, p.adjust < 0.05) differentially expressed genes, respectively. Because phenotypical observations showed that Mp*sdr4* exhibits attenuated responses to ABA-related abiotic stresses, we first examined genes that are not as highly induced by mannitol in Mp*sdr4-2*. To identify such genes, a list of 641 genes significantly down-regulated by at least 1.5 folds (p.adjust < 0.05, log_2_(FC) < (−0.585)) in Mp*sdr4-2* were identified from the transcriptome comparison of mannitol-treated Mp*sdr4-2* and wild-type Tak-1 (Figure 4b). This list was intersected with the genes there were significantly induced by at least 1.5 folds in either Mp*sdr4-2* or Tak-1 (p.adjust < 0.05, log_2_(FC) > 0.585) (Figure 4b). Based on this intersection, we focused on three sets of genes-of-interest: 897 genes that are similarly induced by mannitol in both Tak-1 and Mp*sdr4-2*, and the difference in induction by mannitol is less than 1.5 folds between mutant and wild type (group 897); 245 genes that are more highly induced in Tak-1 by at least 1.5 folds compared to Mp*sdr4-2* (group 245); and 232 genes that are only induced in wild-type Tak-1 (group 232) (Figure 4c). One critical difference between groups 232, 245 with group 897 is that osmotic stress inducibility of genes in the former two groups are strongly or almost completely abolished by the Mp*sdr4* mutation. Because AtSDR4L and its homologs in *Arabidopsis* are transcriptional co-factors, we hypothesize that genes in groups 232, 245 possess different cis-regulatory motifs (CRM) from those in group 897. Motif enrichment analysis supported our hypothesis, and CRMs similar to G-box were identified from both group 232 and group 245, yet no CRM was uncovered from group 897 at an e-value threshold of 10 when DNA sequences 500 base pairs upstream of the transcription start site of genes in these groups were compared to all genes in the *Marchantia* genome by Analysis of Motif Enrichment (AME) (Figure 4d) (McLeay & Bailey, 2010). We further examined specific genes in each of the three groups and visualized their expression using Integrative Genomics Viewer (IGV) (Figure 4e) (Robinson *et al*., 2011). Notable genes in group 232 and 245 include marker gene to osmotic stress (Mp*LAXR*) (Tan *et al*., 2023) and ABA (Mp*MFT*) (Eklund *et al*., 2018), many *LEA* and *LEA-Like* (*LEAL)* genes (e.g. Mp*LEA1*, Mp*DHN1*, Mp*LEAL1*, Mp*LEAL2*) (Jahan *et al*., 2019), and a homolog of ABA biosynthesis gene Mp*NCED* (Tanizawa *et al*., 2025). Interestingly, Mp*ABI3A* is in group 897. Mp*ABI3A* only exhibits modest suppression of induction by mannitol in Mp*sdr4-2*, possibly due to a lack of G-box in its TSS500 sequence. Additionally, we observed that homologs of several cell wall modification genes such as Mp*EXP4* and Mp*PL1.3* are induced by mannitol in Mpsdr4-2 but not in wild-type Tak-1 (Figure 4e), further supporting attenuated inhibition of growth by mannitol in Mp*sdr4* mutants (Figure 3, Figure S4).

**Figure 4.**
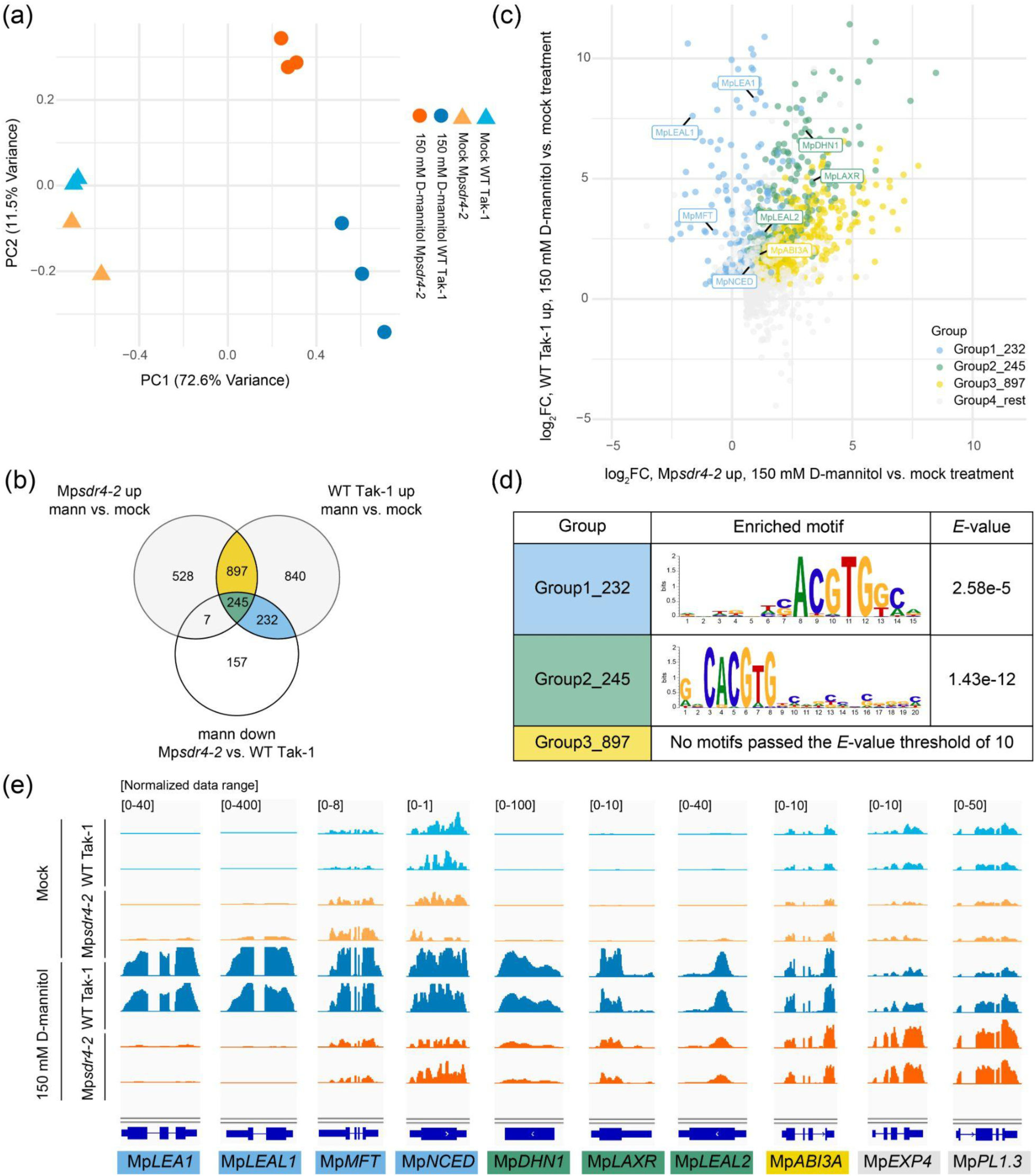
Attenuated ABA response in Mp*sdr4* is linked to the G-box motif. (a) Principal component analysis using multidimensional scaling on the transcriptome of wild-type Tak-1 and Mp*sdr4-2* samples subjected to mock- and 150 mM D-mannitol treatment showed the distance between samples is largely determined by treatment and genotype. (b) Venn diagram to visualize the number of shared and distinct genes that are up-regulated in either or both of wild-type Tak-1 and Mp*sdr4-2* in response to 150 mM D-mannitol (Mann). The top two circles represent genes significantly up-regulated by at least 1.5 folds by D-mannitol in comparison to mock treatment in Mp*sdr4-2* (top left) and Tak-1 (top-right). The bottom circle represents genes significantly down-regulated by at least 1.5 folds in D-mannitol-treated Mp*sdr4-2* in comparison to D-mannitol-treated Tak-1. Yellow, green, and blue colors highlight gene sets that are similarly up-regulated in wild-type Tak-1 and Mp*sdr4-2*, more strongly induced in Tak-1 by at least 1.5 folds, and almost exclusively induced in Tak-1 by D-mannitol, respectively. (c) Satterplot to show log_2_(fold change) of genes that are up-regulated by 1.5 folds in either or both of wild-type Tak-1 and Mp*sdr4-2* in response to 150 mM D-mannitol using the same color code as in (b). (d) Top motifs enriched in both Group 232 and Group 245 include a 5’-ACGTG-3’ core that is similar to G-box. (e) Screenshots of selected genes involved in ABA and osmotic stress responses and cell wall modification visualized in Integrative Genomics Viewer. Two biological replicates are shown for each genotype and treatment. See also Figure S4.

### MpSDR4 inhibits gemma cup formation and growth

In addition to seed development and germination, ABA also influences vegetative to reproductive phase transition in flowering plants. In Arabidopsis, lesions in ABA biosynthesis and signaling often lead to earlier flowering (Shu *et al*., 2018). The reduced ABA sensitivity reflected in thallus growth in Mp*sdr4* prompted us to examine whether the mutant also has faster vegetative to reproductive transition. *Marchantia polymorpha* possesses a gametophyte-dominant life cycle, with the haploid gametophyte also undergoing vegetative reproduction through gemma cup formation. These cups produce gemmae, asexual propagules derived from the cup base (Kato *et al*., 2020). Compared to wild-type Tak-1 and Tak-2, all Mp*sdr4* mutants exhibit earlier formation of gemma cups (Figure 5a). For example, all mutant alleles develop more gemma cups on thalli 20 days after sowing gemmae under long-day conditions, with the most basal cup well-expanded (Figure 5a and 5b). In contrast, growth of gemma cups has just started at this time on wild-type Tak-1, but has not initiated on wild-type Tak-2 thalli (Figure 5b). Furthermore, gemma cups of Mp*sdr4* mutants reach a notably larger average size. At 26 days after sowing under continuous illumination, gemma cups of Mp*sdr4* mutant exhibit a significantly larger average size (*p* < 0.05, one-way ANOVA followed by Tukey’s HSD test) than wild-type Tak-1 (Figure 5c). Cytokinin is essential for gemma cup initiation, while auxin may negatively influence gemma cup formation (Flores-Sandoval *et al*., 2016; Kato *et al*., 2020). An R2R3-MYB transcription factor, GEMMA CUP-ASSOCIATED MYB1 (GCAM1), has been identified as a key regulator for the formation of both gemma cup and gemma (Yasui *et al*., 2019). Mp*GCAM1* is up-regulated in gemma cups and can therefore be used as a marker gene of cup development (Yasui *et al*., 2019). Consistent with the early initiation and overgrowth of gemma cups in Mp*sdr4* mutants, we observed elevated transcript levels of Mp*GCAM1* in Mp*sdr4* mutants in both Tak-1 and Tak-2 backgrounds (Figure 5d).

**Figure 5.**
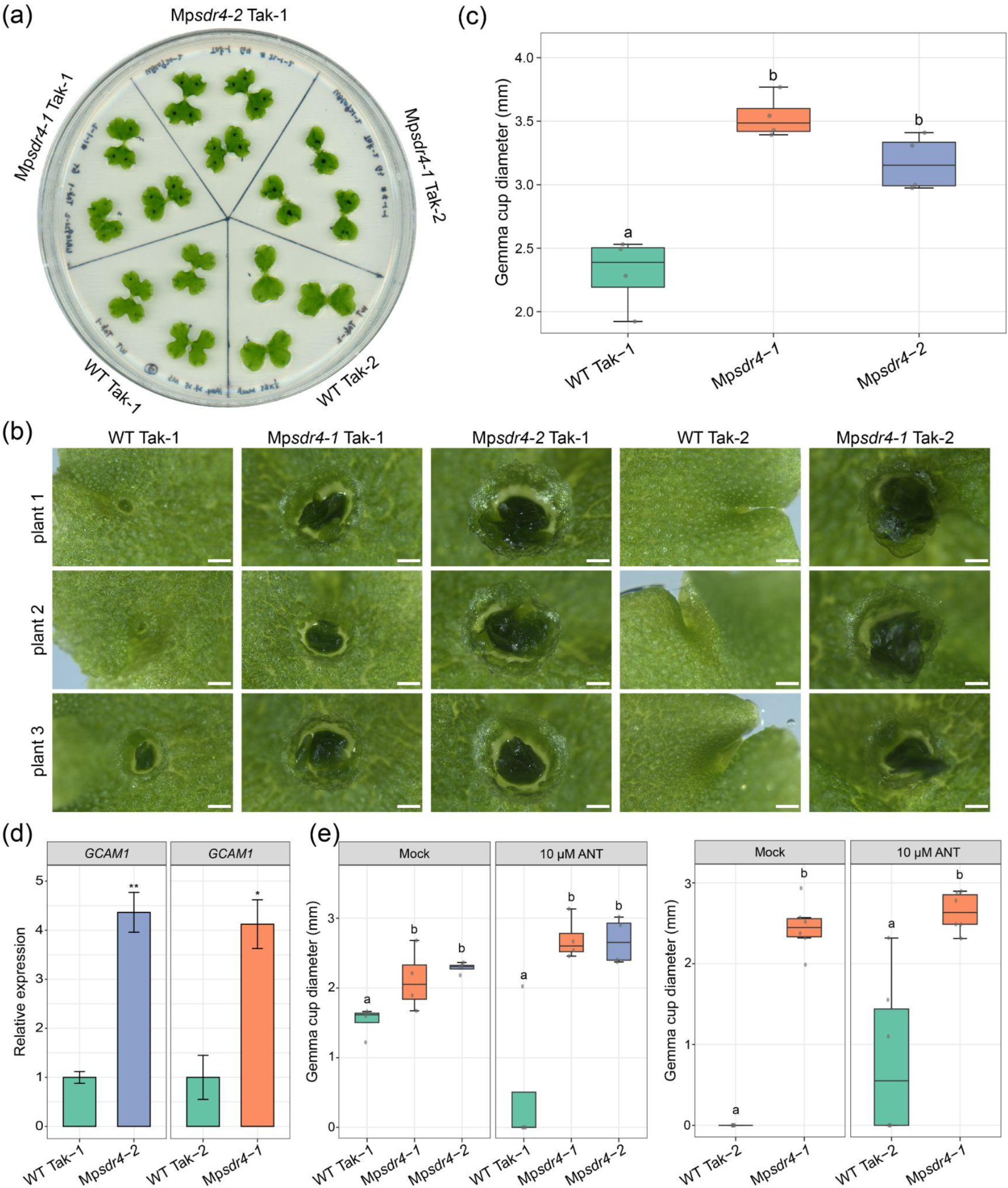
MpSDR4 inhibits gemma cup emergence and expansion. (a - b) Whole plate scanning (a) and magnified images of gemma cups (b) of 20-day-old Mp*sdr4* mutants and the wild type grown on the mock plate under long-day conditions. Scale bars = 1 mm. (c) Gemma cup diameter of 26-day-old Mp*sdr4-1*, Mp*sdr4-2* mutants and wild-type Tak-1 grown under continuous light on mock plates. Different letters represent groups with significant differences (n=4, *p* < 0.05, one-way ANOVA followed by Tukey’s HSD test). (d) RT-qPCR quantification of *GCAM1* transcript abundance from 26-day-old wild-type Tak-1 and Mp*sdr4-2* (left), and 20-day-old wild-type Tak-2 and Mp*sdr4-1* (right). Plants were grown on mock plates under continuous light. *GCAM1* level was normalized to Mp*APT*. Data are displayed as means ± SE (n=3). Significant differences are indicated by asterisks (\**p* < 0.05, \*\**p* < 0.01, two-tailed Student’s *t* test). (e) Gemma cup diameter of 26-day-old Mp*sdr4-1*, Mp*sdr4-2* mutants and wild type in Tak-1 background (left) and Mp*sdr4-1* mutants and wild type in Tak-2 background (right). Plants were grown under continuous light on mock medium and medium supplemented with 10 µM ANT. Different letters represent groups with significant differences. (n=4, *p* < 0.05, one-way ANOVA followed by Tukey’s HSD test; n=6, *p* < 0.05, two-tailed Student’s *t* test). See also Figure S5.

**Figure 6.**
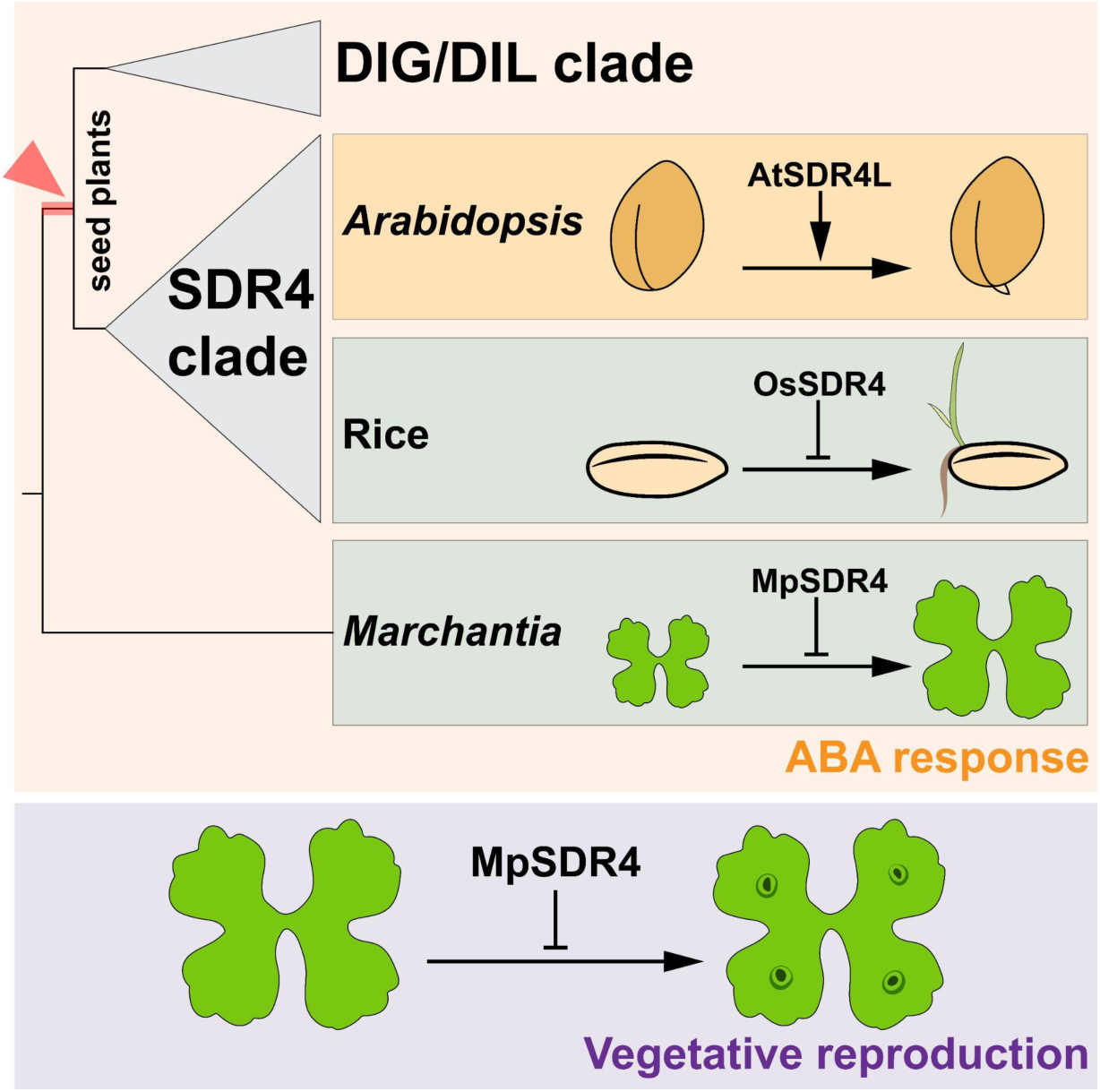
Evolution and functional divergence of SDR4 orthologs in green lineages A model depicting a major duplication event of *SDR4* genes closely preceded with the origin of seed plants, the function of MpSDR4 in ABA response and vegetative reproduction in *Marchantia*, and the functional similarity and divergence of MpSDR4 in comparison to its ortholog in rice and *Arabidopsis*.

We used ABA antagonist antabactin (ANT) (Vaidya *et al*., 2021) to investigate whether the phenotype of gemma cup formation and development in Mp*sdr4* mutants are linked to ABA. We first validated the efficacy of ANT to antagonize ABA in *Marchantia*. In the presence of both ABA and ANT, Mp*sdr4* mutants and wild-type controls grow to a similar size, comparable to those grown on mock treatment plates (Figure S5a). The lack of ABA-induced growth inhibition, and the lack of size difference between Mp*sdr4* mutants and wild type on plates supplemented with both ABA and ANT indicate that ANT can compete efficiently with ABA to block ABA signaling in *M. polymorpha*. We then scored gemma cup growth on medium supplemented with and without ANT. Mp*sdr4* mutants exhibited earlier gemma cup formation and significantly larger gemma cups (*p* < 0.05, one-way ANOVA followed by Tukey’s HSD test for Tak-1 background and two-tailed Student’s *t* test for Tak-2 background) than wild-type Tak-1 and Tak-2 regardless of ANT (Figure 5e and S5b). These results show that MpSDR4 can still regulate gemma cup formation and development when ABA response is blocked, suggesting an ABA-independent pathway for gemma cup development.

## Discussion

### Evolutionary history of the *SDR4* family

Our rooted phylogeny with hypothesis testing supports a paralogous relationship in seed plants between a single SDR4 clade and a single DIG/DIL clade, resulting from a major gene duplication event around the origin of seed plants. This major duplication of *SDR4* and *DIG*/*DIL* was not reported in a previous phylogenetic study that focused solely on *SDR4* genes within angiosperms (Subburaj *et al*., 2016). The timing of the implied duplication is consistent with the ζ event that led to an ancient whole genome duplication in land plants (Jiao *et al*., 2011), and so might coincide with other key innovations in spermatophytes, such as the development of ovules, seeds and pollen (Bowles *et al*., 2020). This coincidence may mark the point at which the evolution of SDR4 became specialized to regulate seed dormancy and germination via ABA signaling (Sun *et al*., 2019). Nearly all seed-plant homologs are found within SDR4 and DIG/DIL clades, indicating the broad retention of both *SDR4* and *DIG*/*DIL* genes following the duplication. A few gymnosperm sequences are associated exclusively with either the SDR4 clade (e.g., Gnetopsida) or the DIG/DIL clade (e.g., *Taxus* and *Pinus*), indicating possible loss of SDR4 or DIG/DIL orthologs in them. The placement of angiosperm homologs in the SDR4 clade is also largely consistent with our overall understanding of angiosperm phylogeny (The Angiosperm Phylogeny Group, 2016). However, the fact that six DIG/DIL homologs of *A. thaliana* are dispersed within the angiosperms DIG/DIL clade, and that some are sister to other eudicot groups that also include dispersed duplicates (e.g., Solanum), supports the idea that there were subsequent rounds of duplication of this gene in eudicots and perhaps other angiosperms. These subsequent duplications may have provided additional raw genetic material for the evolution of novel traits within specific angiosperms lineages. The broad retention of these homologs could have occurred through sub- or neofunctionalization, gene co-option, gene dosage effect, dosage balance, and the rewiring of regulatory networks (Lynch & Conery, 2000; Panchy *et al*., 2016).

### Orthologs of SDR4 are associated with ABA responses and G-box motif in both

#### *Marchantia* and *Arabidopsis*

The core components of ABA signaling, including receptors, phosphatases and kinases, and downstream transcription factors, have already been present in bryophytes (Sun *et al*., 2019). The fact that SDR4 orthologs are also widely present in seed-free lineages and that Mp*sdr4* mutants exhibit altered ABA sensitivity here, suggests that SDR4 might have a very early association in land plants with at least a rudimentary framework of ABA signaling. Consistent with the faster growth of Mp*sdr4* mutants than wild-type *Marchantia* on ABA, mannitol, and salt plates, transcriptome analysis showed that a large subset of stress responsive genes are up-regulated to a lower level in Mp*sdr4* plants than wild type (Figure 4c). More interestingly, the genes that showed suppressed induction are enriched with G-box (5’-CACGTG-3’) in their promoters (Figure 4d). Thus, we have established a link between SDR4 and G-box in *M. polymorpha*. Notably, G-box is also enriched in differentially expressed genes of At*sdr4l* mutant seedlings in *Arabidopsis* (Wu *et al*., 2022). The main difference between *Arabidopsis* and *Marchantia* is that the G-box associated DE genes in At*sdr4l* mutant seedlings are highly up-regulated compared to wild-type *Arabidopsis*, in contrast to a weaker induction by osmotic stress in Mp*sdr4* than in wild-type *Marchanita*. These molecular differences of G-box associated genes are in line with elevated and reduced sensitivity to ABA and osmotic stress of At*sdr4l* and Mp*sdr4* mutants, respectively.

At the level of individual marker genes, the reduced sensitivity to ABA of Mp*sdr4* mutants in thallus growth is corroborated by a drastic reduction in the induction ABA- and osmotic stress-responsive genes such as Mp*LEA*s, Mp*LEAL*s, Mp*MFT*, and Mp*LAXR* (Figure 4e). The orthologs of *LEA*s and *MFT* are known to be ABA-responsive in a broad range of seed and seed-free species (Xi *et al*., 2010; Eklund *et al*., 2018; Artur *et al*., 2019; Song *et al*., 2020). Altered inductivity of these genes in response to osmotic stress in Mp*sdr4* supports at the molecular level the possibility that *SDR4* family was a component of ABA responses in early land plants. The B3 transcription factor ABI3 is a core component in ABA signaling and is functionally conserved in ABA responses in green lineages. Orthologs of *ABI3* from moss and maize can complement *abi3* mutants in *Arabidopsis* (Suzuki *et al*., 2001; Marella *et al*., 2006), and MpABI3A restores the expression of an ABA reporter gene in the Pp*abi3abc* triple mutant in *P. patens* (Eklund *et al*., 2018). Although the induction of Mp*ABI3A* in mannitol-treated Mp*sdr4-2* is not strongly attenuated compared to that in mannitol-treated Tak-1 (Figure 4c), the difference is still significant (log_2_(FC) = (−0.44), p.adjust = 0.012). This difference in induction can also be viewed by the comparisons between mannitol and mock treated transcriptomes of the same genotype. Mp*ABI3A* is more weakly induced by mannitol in Mp*sdr4-2* (log_2_(FC) = 1.10, p.adjust = 1.6E-04) than in Tak-1 (log_2_(FC) = 1.81, p.adjust = 2.34E-05) (Figure 4e). Therefore, MpABI3A may also contribute to MpSDR4-mediated modulation of ABA sensitivity.

The rudimentary association of *SDR4* family with ABA signaling could have subsequently become tailored for the regulation of seed maturation and acquisition of dormancy in spermatophytes. Seed dormancy is an adaptive trait that helps avoid germination in adverse environments (Finch-Savage & Footitt, 2017). Among the major types of seed dormancy, physiological dormancy (PD) is prevalent in most angiosperm lineages, including the monocots and eudicots (Willis *et al*., 2014). PD is maintained by ABA, and break of dormancy by physiological conditions often results in a reduction in ABA level and sensitivity (Weitbrecht *et al*., 2011). Mutants of SDR4 in rice and Arabidopsis exhibit opposite germination behavior and sensitivity to ABA and related stresses. In rice, Os*sdr*4 mutant germinates more readily than wild type (Chen *et al*., 2021; Zhao *et al*., 2022). By contrast, *Arabidopsis* At*sdr4l* exhibits enhanced dormancy (Cao *et al*., 2020) and elevated sensitivity to osmotic stress (Lu *et al*., in revision). Because vegetative growth of Mp*sdr4* mutants is less sensitive to ABA and osmostress, enhancing ABA sensitivity might be a primitive function (symplesiomorphy) of SDR4. Alternatively, SDR4 might have evolved to execute lineage-specific regulation of ABA sensitivity. In flowering plants, many genes involved in seed maturation possess both G-box and RY (5’-CATGCA-3’) motifs (Nakabayashi *et al*., 2005; Jo *et al*., 2020), and the physical interaction of SDR4 with B3 TF VIVIPAROUS1/ABI3-LIKE2 (VAL2) could be one mechanism that recruits SDR4 close to a G-box motif in *Arabidopsis* (Lu *et al*., 2024; Go *et al*., 2024). An interesting future direction is to examine the mechanism that links G-box-containing genes and MpSDR4 in *Marchantia*.

## Conclusion

In summary, our study revealed a key duplication event in the molecular evolution of the SDR4 family. Functional characterization of MpSDR4 suggests that the association of SDR4 ortholog and ABA responses predates the evolution of spermatophytes. Further investigations are needed to understand the more mechanistic underpinnings how MpSDR4 interacts with other regulators of ABA responses and gemma cup growth, and to explore the biological basis for the retention and differentiation of both the SDR4 clade and the paralogous DIG/DIL clade in seed lineages, which may help shape distinct patterns of regulation of dormancy in rice and Arabidopsis. We will gain more insights into hormonal and transcriptional regulation by studying families like SDR4 that show ABA-related yet lineage-dependent roles, in addition and in contrast to transcription factors such as ABI3 that have highly conserved functions across land plants (Khandelwal *et al*., 2010; Eklund *et al*., 2018).

## Supplemental information

● Supplemental Table 1. Compiled sequence for SDR4 protein family phylogenetic reconstruction

● Supplemental Table 2. Oligonucleotides (5’ - 3’) for cloning, genotyping, and RT-qPCR

● Supplemental Figure 1. Sequence alignment of SDR4 homologs

● Supplemental Figure 2. Cladogram of the constrained tree search output for the SDR4 protein family

● Supplemental Figure 3. Mp*sdr4* mutants are hyposensitive to a broad range of ABA concentrations

● Supplemental Figure 4. Mp*sdr4* mutants in Tak-2 background are hyposensitive to osmostress

● Supplemental Figure 5. Antabactin antagonizes ABA response in Marchantia but does not inhibit elevated growth of gemma cup in Mp*sdr*4 mutants

## Acknowledgements

We thank Drs. Takayuki Kohchi (Kyoto University) and Tomokazu Kawashima (University of Kentucky) for sharing wild-type Tak-1 and Tak-2, Dr. Cătălin Voiniciuc (University of Florida) and Djamei Lab (University of Bonn) for sharing the *Marchantia* transformation protocols, Dr. John Bowman (Monash University) for general discussions of *Marchantia*, Drs. Mark Estelle (UC San Diego) and Michael Prigge (UC San Diego) for discussions about the project, Dr. Mary Berbee and UBC Department of Botany for research awards to support Jiayi Shan’s Honours project, and Bailan Lu for various technical support.

## Author contributions

Conceptualization: L.S. J.S.; Formal analysis: J.S., J.K., L.S.; Funding acquisition: L.S.; Investigation J.S., J.K., L.S.; Resources: J.W., S.C.; Supervision: L.S., S.G.; Validation; J.S., J.K.; Visualization J.S., J.K., L.S.; Writing – original draft: J.S., L.S., J.K.; Writing – review & editing: L.S., J.S., S.G.

