## Supplemental Table 2 and supplemental figures for "Liverwort MpSDR4 regulates abscisic acid sensitivity and gemma cup growth"

### Supplemental Information

#### SUPPLEMENTAL TABLES

**Supplemental Table 2: Oligonucleotides (5’ - 3’) for cloning, genotyping, and RT-qPCR.**

| Mp2g17570_gRNA1_F | CTCGGAGATGAGAGCTGACCATGGGT |
| --- | --- |
| Mp2g17570_gRNA1_R | TAAAACCCATGGTCAGCTCTCATCTC |
| Mp2g17570_gRNA2_F | CTCGGTGCAAGAATCTGTTATCATGT |
| Mp2g17570_gRNA2_R | TAAAACATGATAACAGATTCTTGCAC |
| Mp2g17570_gRNA3_F | CTCGCCAGTCTTGCCAGTGGGCTTGT |
| Mp2g17570_gRNA3_R | TAAAACAAGCCCACTGGCAAGACTGG |
| Mp2g17570_gRNA4_F | CTCGGTACCGCGAGAGTATGTGCTGT |
| Mp2g17570_gRNA4_R | TAAAACAGCACATACTCTCGCGGTAC |
| Mp2g17570_GT_F | ACACAGCGCCCCGAACTCCTAT |
| Mp2g17570_GT_R | GGAGCTCGCTTTGGCACTGGTT |
| pC_F (Sauret-Güeto et al., 2020) | GCAACGCTCTGTCATCGTTAC |
| pC_R (Sauret-Güeto *et al.*, 2020) | GTAACTTAGGACTTGTGCGACATGTC |
| pCsA_MpEF1a_seq | AGTTACCCAAATATCGATACCCGT |
| pCsA_HygR_seq | ACGCAATCGTCCGATCCGGA |
| pCsA_MpU6_seq_F | ACAGTTGAGACACGTGGCGCTG |
| pCsA_MpU6_seq_R | GCTGAGTTTGCGATGTTTTGAGC |
| Mp3g25140_MpAPT_q1F | CGGTGAACGCTGTTTGATTG |
| Mp3g25140_MpAPT_q1R | CGCGCTCTAGTAGCCTTATTC |
| Mp5g23710_MpLEA1_q1F | CTCTACCGAGGCCAACATTATAG |
| Mp5g23710_MpLEA1_q1R | GAGTCTGGAAACGATCTACACAA |
| Mp6g07540_MpDHN1_q1F | GCAACACCAACTCCCAATATG |
| Mp6g07540_MpDHN1_q1R | CCCGGAGTGATCTTGTCTTT |
| Mp6g04830_MpGCAM1_q1F | CTGTACCTTTCAGCGGAGATTG |
| Mp6g04830_MpGCAM1_q1R | GGATTAGTAGCAGCCGCAATAG |

#### SUPPLEMENTAL FIGURES

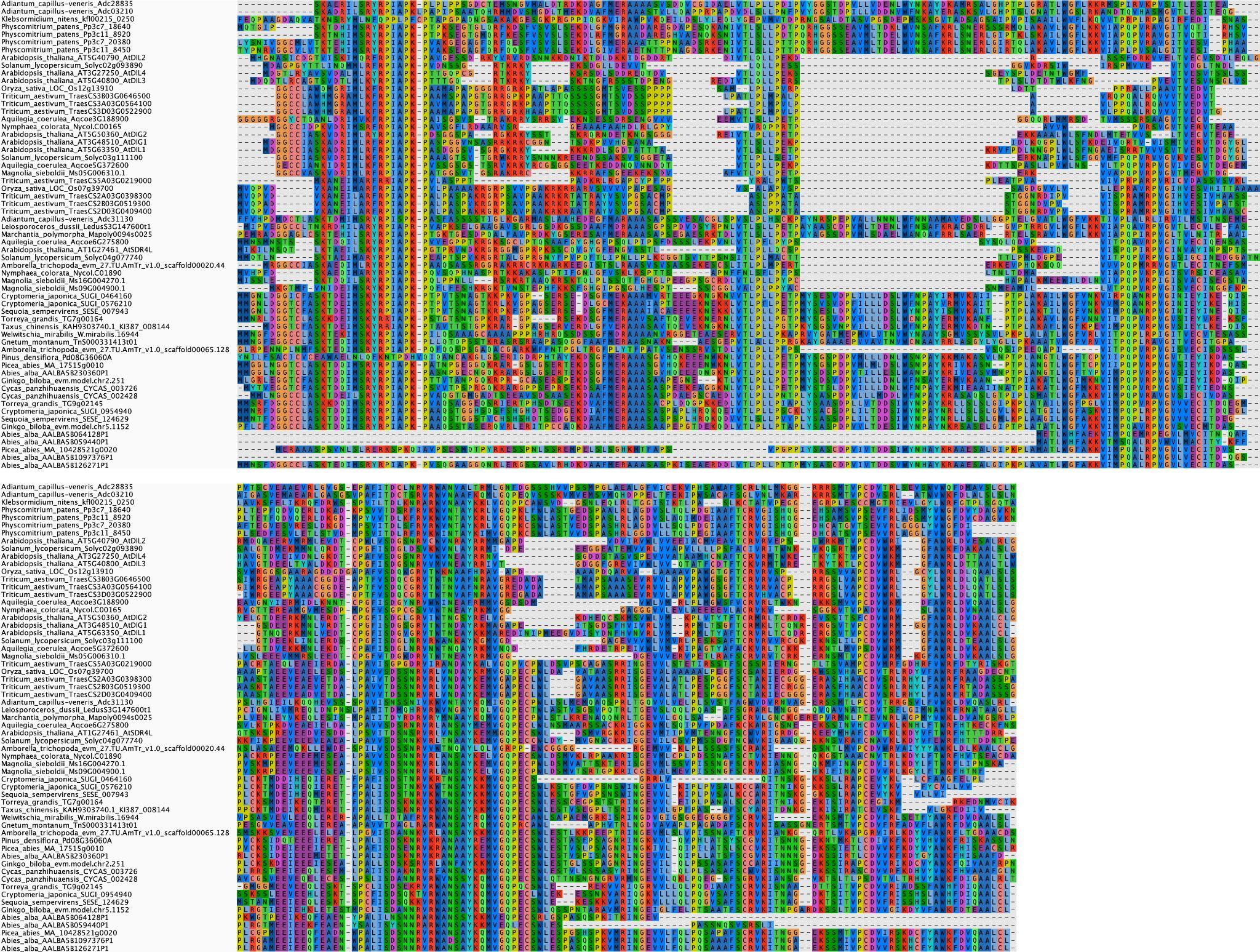

**Figure S1. Sequence alignment of SDR4 homologs**

Trimmed multiple protein sequence alignment of SDR4 protein family for maximum-likelihood (ML) analysis. The multi-sequence alignment of 24 species was trimmed with trimAl and visualized on Aliview.

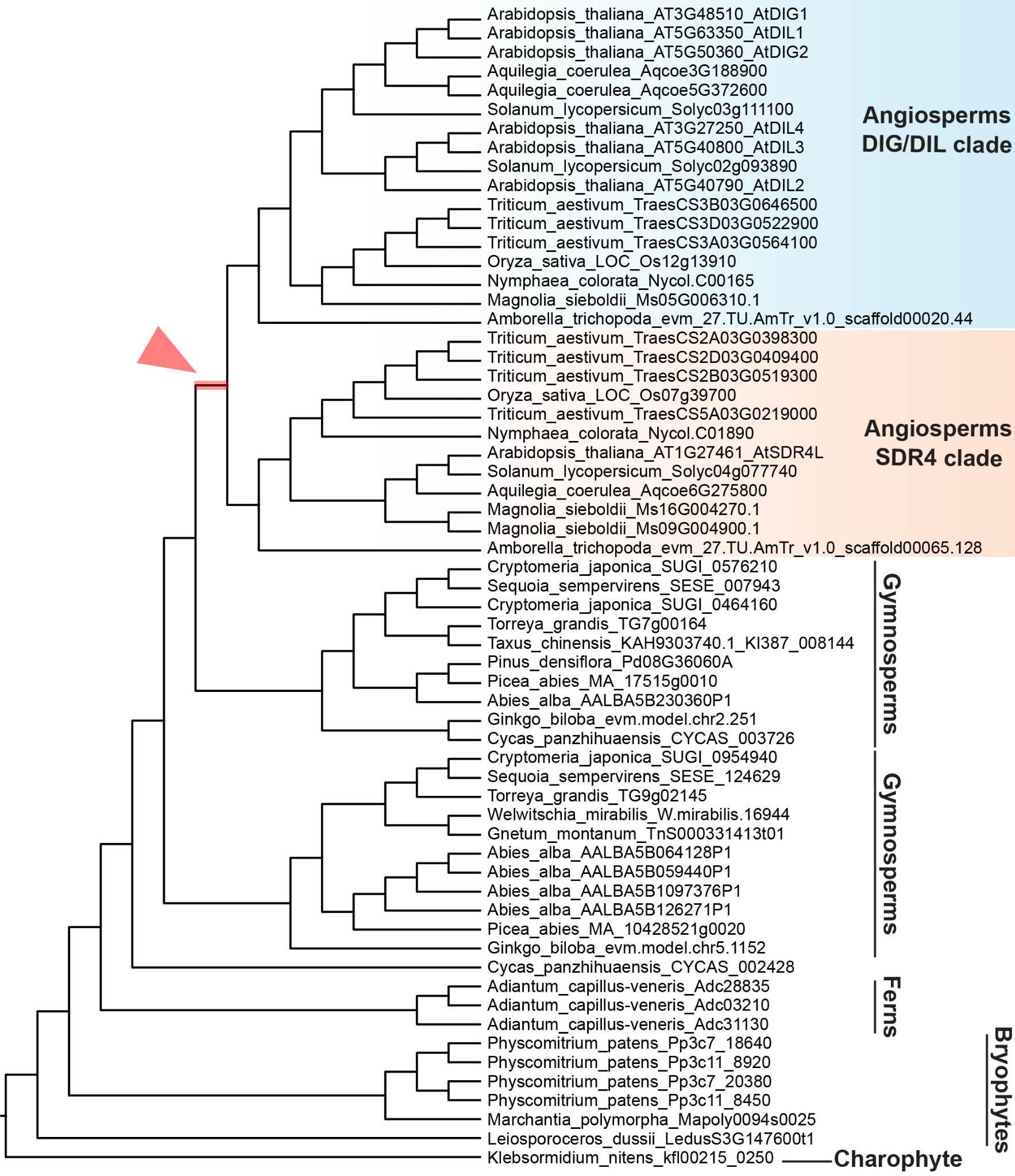

**Figure S2. Cladogram of the constrained tree search output for the SDR4 protein family**

The angiosperms SDR4 clade and angiosperm DIG/DIL clade form a monophyletic group. The red-highlighted branch marked by the arrowhead indicates a possible placement of a gene duplication event around the origin of angiosperms. This tree topology represents an alternative hypothesis (H_a_) tested using the AU test.

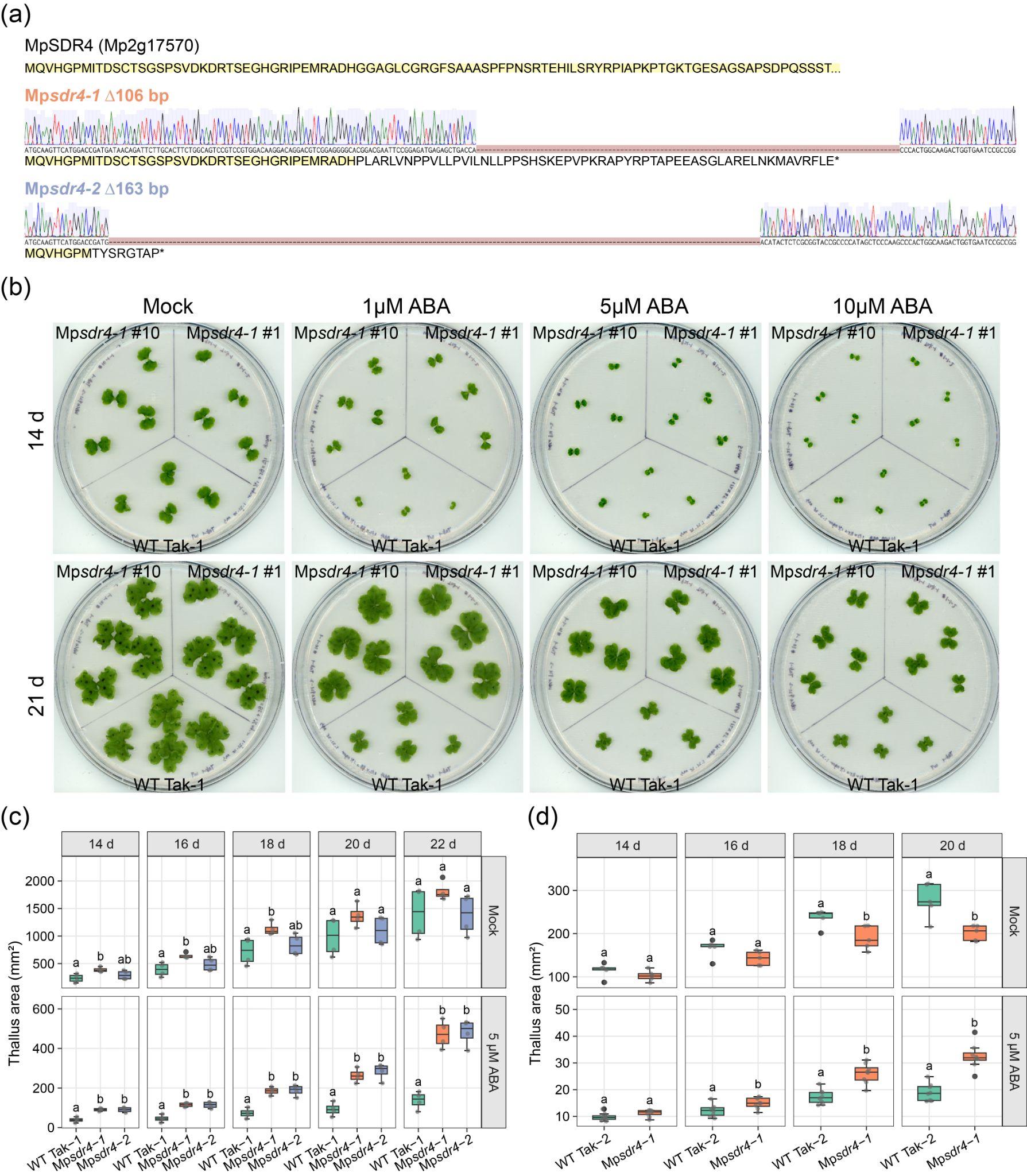

**Figure S3. Mp*sdr4* mutants are hyposensitive to a broad range of ABA concentrations**

(a) Protein sequences of wild-type and mutant MpSDR4. Chromatograms show the DNA sequence of indicated mutant alleles and the segmental deletions. The wild-type protein sequence is highlighted in yellow.

(b) 14- and 21-day-old Mp*sdr4-1* mutants and the wild-type in the Tak-1 background, grown under continuous light on the mock plate and plates containing a serial ABA concentration. Mp*sdr4-1* #10 carries the same mutation as Mp*sdr4-1* #1 but is descended from an independent transformant. All other Mp*sdr4-1* mutants used in this paper were derived from the #1 line.

(c) Thallus area of wild-type Tak-1 and Mp*sdr4* mutants measured at 2-day intervals from day 14 to day 22. Plants were grown under long-day conditions on medium without (mock) or with 5 µM ABA. Different letters represent significant differences (n = 4, *p* < 0.05, one-way ANOVA followed by Tukey’s HSD test).

(d) Thallus area of wild-type Tak-2 and Mp*sdr4* mutants measured at 2-day intervals from day 14 to day 20. Plants were grown under long-day conditions on medium without (mock) or with 5 µM ABA. Different letters represent significant differences (n = 5 for mock and n = 8 for 5 µM ABA treatment, *p* < 0.05, two-tailed Student’s *t* test).

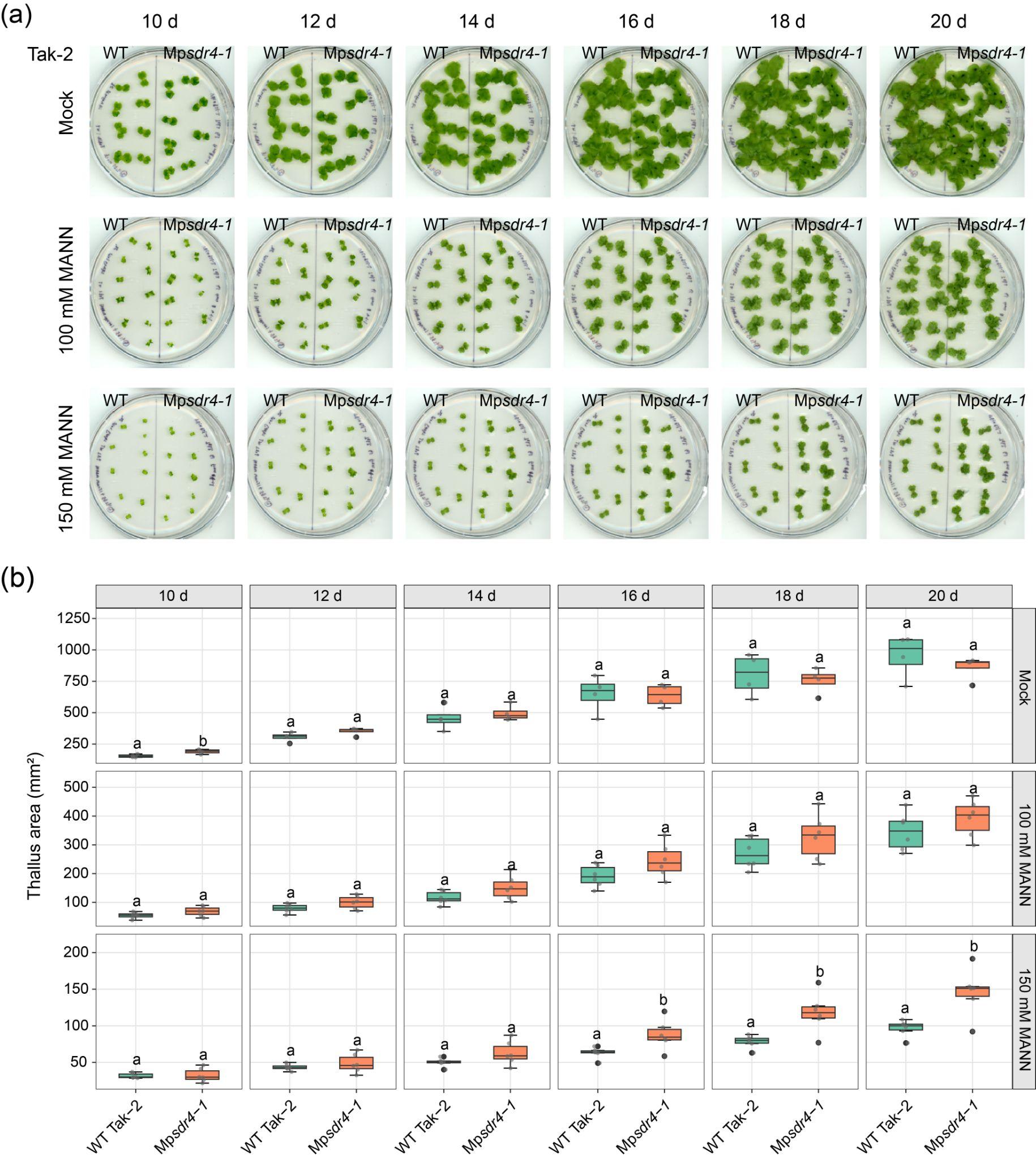

**Figure S4. Mp*sdr4* mutants in Tak-2 background are hyposensitive to osmostress**

(a - b) Scanned plates (a) and size quantification (b) of wild-type Tak-2 and *Mpsdr4* mutant grown on mock medium or medium supplemented with 100 mM and 150 mM D-mannitol. Different letters represent significant differences (n = 4 for mock and n = 6 for mannitol treatment, *p* < 0.05, two-tailed Student’s *t* test).

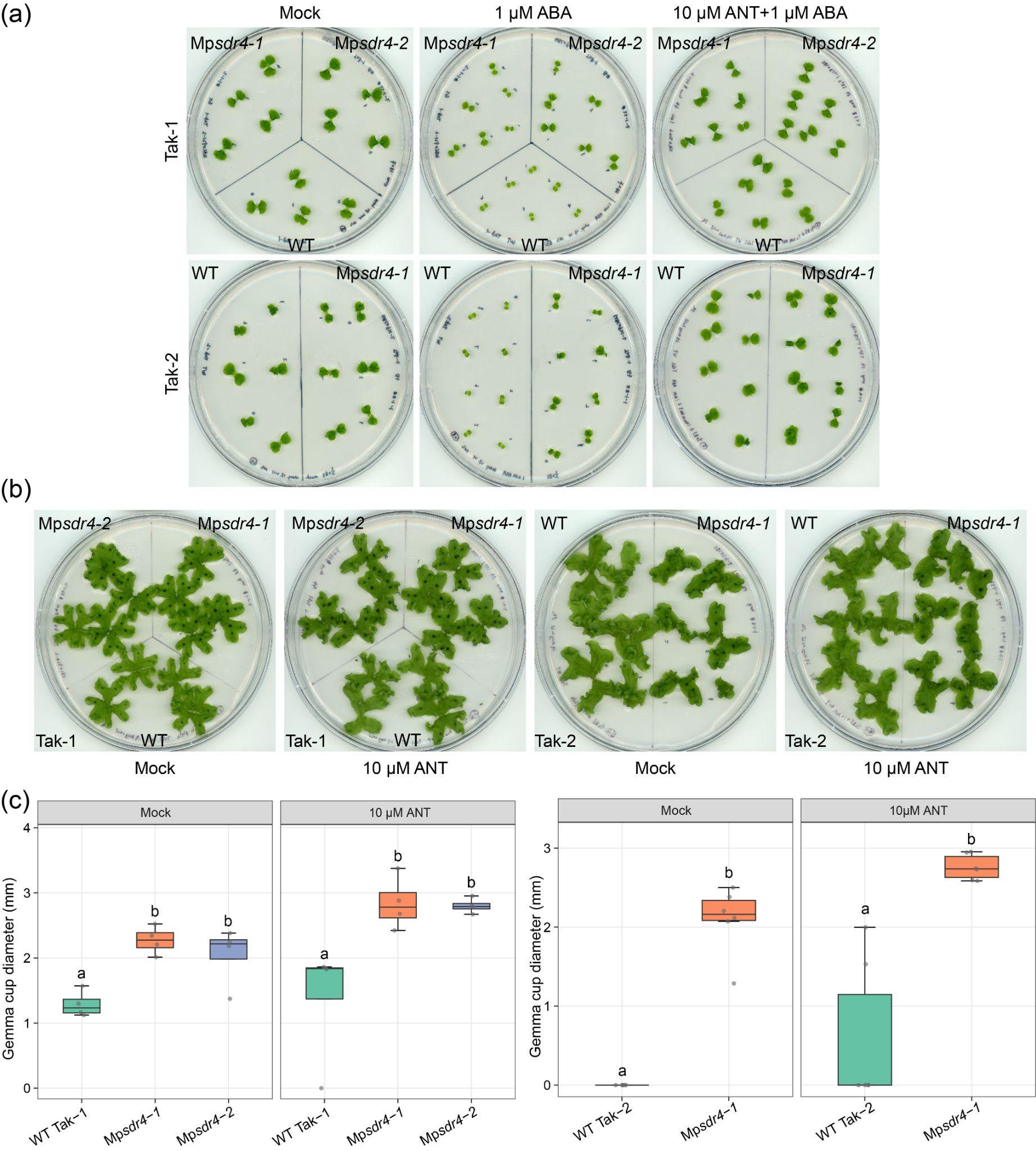

**Figure S5. Antabactin antagonizes ABA response in Marchantia but does not inhibit elevated growth of gemma cup in Mp*sdr4* mutants**

(a) Plate scanning of 16-day-old Mp*sdr4* mutants and the wild-type Tak-1 and Tak-2 grown under long day conditions on mock, 1 µM ABA, and 1 µM ABA plus 10 µM ANT plates.

(b - c) Plate scanning (b) and measurement of gemma cup diameter (c) of 26-day-old Mp*sdr4-1*, Mp*sdr4-2* mutants and wild type in Tak-1 background (left) and Mp*sdr4-1* mutants and wild type in Tak-2 background (right). Plants were grown under continuous light on mock medium and medium supplemented with 10 µM ANT. Different letters represent groups with significant differences. (n=4, *p* < 0.05, one-way ANOVA followed by Tukey’s HSD test; n=6, *p* < 0.05, two-tailed Student’s *t* test). These plates are additional replicates to data shown in Figure 5e.
